# Quantifying the bioavailable energy in an ancient hydrothermal vent on Mars and a modern Earth-based analogue

**DOI:** 10.1101/2022.09.09.507253

**Authors:** Holly R. Rucker, Tucker D. Ely, Douglas E. LaRowe, Donato Giovannelli, Roy E. Price

## Abstract

Putative alkaline hydrothermal systems on Noachian Mars were potentially habitable environments for microorganisms. However, the types of reactions that could have fueled microbial life in such systems and the amount of energy available from them have not been quantitatively constrained. In this study, we use thermodynamic modeling to calculate which catabolic reactions could have supported ancient life in a saponite-precipitating hydrothermal vent system in the Eridania basin on Mars. To further evaluate what this could mean for microbial life, we evaluated the energy potential of an analogue site in Iceland, the Strytan Hydrothermal Field (SHF). Results show that out of the 85 relevant redox reactions that were considered, the highest energy-yielding reactions in the Eridania hydrothermal system were dominated by methane formation. By contrast, Gibbs energy calculations carried out for Strytan indicate that the most energetically favorable reactions are CO_2_ and O_2_ reduction coupled to H_2_ oxidation. In particular, our calculations indicate that an ancient hydrothermal system within the Eridania basin could have been a habitable environment for methanogens using NH_4_^+^ as an electron acceptor. Differences in Gibbs energies between the two systems were largely determined by oxygen – its presence on Earth and absence on Mars. However, Strytan can serve as a useful analogue for Eridania when studying methane producing reactions that do not involve O_2_.

## 1. Introduction

From an astrobiological perspective, a habitable environment can be defined as any setting that can support the metabolic activity of at least one organism (Cockell et al., 2016; Merino et al., 2019). This definition can be translated to places that have enough energy, water and nutrients to sustain life. For many nearby planetary bodies, a sufficient combination of these variables likely could only exist in the subsurface. In particular, it has been speculated that subsurface hydrothermal vents (Shock, 1997; Varnes et al., 2003; Marlow et al., 2014) could host life elsewhere since those on Earth today support large and diverse microbial populations that take advantage of the disequilibria that occurs when reduced, electron donor-rich hydrothermal fluids mix with oxidant-rich seawater. If hydrothermal vents on ancient Mars were also places where redox chemical disequilibria could be sustained by similar fluid mixing, they could have been critical zones for hosting life. Although a number of studies have quantified the amount of energy available for microbial life in hydrothermal systems on Earth (see Lu et al., 2020), determining this for hydrothermal systems on Mars requires knowledge of the rock types and fluids that form hydrothermal systems.

As life originated and began to evolve on Earth during the late Hadean and early Archean eons (4.1-3.5 Gya), Mars entered the Noachian period, a time characterized by geological activity, volcanism, and liquid water (Marlow et al., 2014). Approximately 3.8 billion years ago, Mars is thought to have once been an “ocean world”, which is defined as any planetary body that has, or once had, a liquid ocean, with evidence of fluvial erosion from ancient, dendritic valley networks, putative ocean shorelines, and crater lakes (Wordsworth, 2016; Hendrix et al., 2019). Evidence of volcanic interaction with surficial and subsurface water suggests that hydrothermal systems were likely present on Mars during this period (Griffith et al., 1997; Marlow et al., 2014). Phyllosilicate clay deposits are frequently found as alteration products or direct precipitates in hydrothermal systems on Earth (Cuadros et al., 2013), and it has been suggested that many of the abundant Noachian phyllosilicate clay deposits found on Mars could be of hydrothermal origin (Mustard et al., 2008; Michalski et al., 2017; Michalski et al., 2018).

The Eridania basin, located within the southern highlands on Mars, is thought to have had seafloor hydrothermal activity during the Noachian-Early Hesperian periods (approximately 4.0 to 3.5 Gya) (Michalski et al., 2018). It likely held an ancient inland sea approximately a million km^2^ in size with a depth range of 1-1.5 km (Michalski et al., 2017; Irwin et al., 2004; Pajola et al., 2016), resulting in a sea with a volume greater than that of all other basin lakes on Mars and nearly ten times the volume of the Great Lakes along the US-Canadian border (Michalski et al., 2017). Eridania is also host to the strongest remaining evidence of magnetism on Mars and has been proposed to have been a crustal spreading region in the past, similar to Earth’s mid-ocean ridge spreading centers (Michalski et al., 2017). Finally, the ratio of volume to watershed area (V:A_w_) of Eridania is anomalously high (102 m), which suggests that the sea had a groundwater source rather than precipitation (Fassett et al., 2008).

Many deposits preserved in the oldest rocks on Mars represent materials that were seemingly exhumed from the subsurface (Ehlmann et al., 2011). For example, the Compact Reconnaissance Imaging Spectrometer for Mars (CRISM) has revealed Fe/Mg clay minerals such as saponite, talc, serpentine, sepiolite, and nontronites in Eridania Basin. (Michalski et al., 2017). The presence of these Fe/Mg-phyllosilicates suggests that the minerals precipitated in an alkaline environment since layered phyllosilicates, such as saponite and smectite, tend to form under high pH (Adeli et al., 2015). Additionally, long-term water-rock interactions are important for the formation of clays, further supporting the notion that an established sea existed in this region (Ehlmann et al., 2012).

Since the inaccessibility of putative ancient hydrothermal systems on Mars prevents a thorough investigation of their habitability, we turn to similar geochemical environments on Earth to better constrain their biological potential. Icelandic rocks and hydrothermal systems are a convenient analogue for Noachian basalts and basaltic alteration (e.g., Allen et al, 1981; Ehlmann et al., 2012; Black et al., 2018). In particular, the vent chimneys found in the Strytan Hydrothermal Field (SHF), located in Eyjafjord, (Eyjafjörður in Icelandic), are made of saponite and emit alkaline fluids (end-member pH ∼10) at low temperature (70 °C) and are enriched in dissolved silica (Marteinsson et al., 2001; Geptner et al., 2002). SHF is a shallow-sea hydrothermal vent field (16 to 70 m below sea level) consisting of several tall cone-like chimneys (up to 55 m high monoliths) (Marteinsson et al., 2001; Price et al., 2017). Unlike many basalt-hosted submarine hydrothermal systems, the hydrothermal fluid is derived from meteoric-originated groundwater (Marteinsson et al., 2001). Another unusual characteristic of Strytan fluids is that the high pH is not primarily due to serpentinization (McCollom, 2007; Allen et al., 2004), but also hydrogen ion metasomatism, in which protons are exchanged for cations within the rock (Price et al., 2017). Additionally, calcium carbonate (CaCO_3_) precipitation in the rocks exhausts the already limited carbon dioxide (CO_2_) buffer from its closed-system groundwater source, further allowing for the increase of the pH of the hydrothermal fluids (Price et al., 2017). As a shallow-sea, basalt-hosted vent field with saponite precipitation from an alkaline hydrothermal fluid, SHF is a strong analogue for an Eridania hydrothermal system. One way to compare these sites is to investigate their energetic potential. Energetic analyses have been successfully executed in the study of many hydrothermal vents, including deep-sea vents (e.g., McCollom et al., 1997; Shock et al., 2005; Amend et al., 2011), shallow-sea vents (e.g., Akerman et al., 2011; Price et al., 2015; Lu et al., 2020), and terrestrial systems (e.g., Inskeep et al., 2005; Spear et al., 2005; Shock et al., 2010). Energetics calculations are a powerful tool for analyzing potential microbial metabolisms and investigating habitability using a quantitative point of reference (Hoehler et al., 2007). The overall objective of this study was to use energetics calculations to determine the potential habitability of a saponite-precipitating hydrothermal system on ancient Mars, while also comparing it to the SHF, evaluating its usefulness as a Mars analogue. To study the energetics of Eridania and SHF, measured (SHF) and modeled (SHF and Eridania) hydrothermal fluids, created based on the environmental characteristics of each vent system, were used. Modeling the Strytan hydrothermal fluids is crucial, as it provides a ground-truth for the modeling conducted for the Eridania fluids. Energetics calculations were then performed for metabolically relevant chemical reactions to determine the most favorable metabolisms at both SHF and Eridania and to compare the two vent systems.

## 2. Materials and Methods

### 2.1 Modeling the Noachian Eridania hydrothermal system

#### 2.1.1 Eridania Rainwater

The concentrations of the *i*th chemical species, *C*_*i*_, in Eridania rainwater were calculated from the solubility of gases in an assumed Noachian atmosphere using

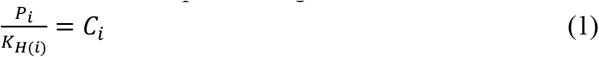

where *P*_*i*_ stands for the partial pressure of the *i*th gas and *K*_*H (i)*_ denotes the Henry’s Law constant for that gas. The partial pressure of ancient Mars used in this study are listed in Table 2, with a total pressure of 1.64 bars. Values of *K*_*H (i)*_ at 273 K were calculated using

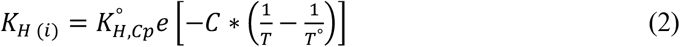

where, *K*º_*H,Cp*_ is the standard Henry’s law constant for a gas at 298 K (Sander, 2015), *C* is a constant (Sander, 2015), *T* = 273 K and *T*º = 298 K. The gases were then equilibrated with pure water at pH 4, to represent an acidic Noachian rainwater (Catling, 1999). The dissolved gases were also speciated in the rainwater at 2 ºC, *P* = 1 bar and 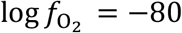, consistent with values used in similar studies (Varnes et al., 2003).

The atmospheric model represents a period of warm, reducing conditions during the atmospheric fluctuations of Noachian Mars (Wordsworth et al., 2021). A partial pressure of 1.5 bar for CO_2_ at 273 K was chosen such that the surface of Mars would have been warm enough for liquid water to exist (Wordsworth, 2016; Wordsworth et al., 2017). The next most abundant gases in this model are CH_4_ and H_2_, both at 3.5% of the atmosphere (Wordsworth et al., 2017; Table 2). The inclusion of carbon monoxide in the model is important as it allows for speciation of additional carbon-bearing species. The concentration of CO was estimated based on modern Mars values (0.007 %) due to the lack of this trace gas in Noachian atmospheric models (Owen et al., 1977). Nitrous oxide (NO) is also needed for speciation of nitrogen-bearing species, and the amount of NO in the atmospheric model was estimated at 0.1 % to represent a trace gas.

Noble gases argon (Ar), helium (He), and neon (Ne), in addition to nitrogen (N_2_) were set at 0.25 %. Sulfur dioxide (SO_2_), an outgassed product of volcanism, was likely present in the atmosphere but is currently not well constrained in Noachian climate models. The current upper limit of SO_2_ in the Martian atmosphere (0.3 ppb; Encrenaz et al., 2011) was used as a constraint for total sulfur in the atmosphere. A conservative estimate of HCl in the atmosphere (0.2 ppb) was used based on current atmospheric conditions (Villanueva et al., 2013). Presently, Noachian atmospheric models are not robust enough to include estimations for more trace gases, so modern Mars values were used where needed.

#### 2.1.2 Modeled Mineralogy of Eridania

The rock types used to calculate the consequences of water-rock interactions on Mars are based on the composition of basalt found north of Eridania, the Backstay rock. The Backstay rock has been directly observed by a rover and therefore has a known oxide composition (McSween et al., 2006; Squyres et al., 2006). It was found in the Columbia Hills of the Gusev Crater (Squyres et al., 2006), which is connected to the Eridania basin by Ma’adim Vallis, a large outflow river valley that originates in Eridania (Irwin et al., 2004). Although the Backstay rock contains 11-16% olivine (McSween et al., 2006; Squyres et al., 2006), we considered two water-rock scenarios in which the total olivine content of the rock was set to 5 and 16%, thereby including other potential rocks in the system that might be low in olivine. The olivine-rich sample uses the major oxide content from McSween et al. (2006) and the proportion of forsterite to fayalite in the 16 wt % olivine sample (0.54 and 0.46, respectively) was used to calculate the amount of each endmember in the 5 wt % olivine model. The proportion of Fe^III^ (and therefore Fe_2_O_3_) to total iron was kept at 0.23, as calculated in the 16 wt % sample (McSween et al., 2006). The amount of forsterite and fayalite was used to calculate the changes in total SiO_2_, MgO, FeO, and Fe_2_O_3_ content of Backstay rock, as these oxides would be directly affected by varying the olivine wt % (Table 3). The content of the other major oxides (e.g., TiO_2_, Al_2_O_3_, MnO, MgO, CaO, Na_2_O, K_2_O, and P_2_O_5_) from the 16 % olivine sample were used for both samples (McSween et al., 2006). The total pressure at which the water-rock reactions were conducted followed the same calculation as Iceland (see below). Using the gravity of Mars, the total pressure was calculated to be 28 bar. The calculations were carried out from 0 to 140 ºC, but the fluid output at 72 ºC was chosen to compare the energetics at the same temperature as Strytan.

### 2.2 Modeling Strytan hydrothermal fluids

#### 2.2.1 Strytan water

Icelandic rainwater at 7^°^C (Arnorsson and Andresdottir, 1995) was used as the basis for the composition of the fluid that was reacted with rocks at Strytan (see Supplemental Table 1), in line with observations that meteoric water feeds the hydrothermal vents there (Price et al., 2017). Calculations were carried out at 78 bars, assuming that the reactions were taking place 200 m beneath basalt and under the tallest saponite cone at SHF (55 m) and the local water depth. The depth of 200 m was used to represent a shallow entrainment of meteoric originated ground water into the subsurface during hydrothermal fluid evolution (McMullin et al., 2000). The densities of saponite and basalt used were 2.3 kg/m^3^ and 2.9 kg/m^3^, respectively (Christensen et al., 1982; Anthony et al., 2001). Similar to Eridania Basin, calculations were carried out from 0-150 °C to encompass the reservoir temperature at SHF (50-100 ºC) (Geptner et al., 2002).

**Table 1.** Elemental abundances of Icelandic olivine-basalt rocks.

| Element | Molality (mol/kg) |
| --- | --- |
| Si | 7.958 |
| Ti | 0.220 |
| Fe | 1.638 |
| Mg | 1.776 |
| Al | 2.796 |
| Ca | 2.034 |
| Na | 0.752 |
| K | 0.042 |
| P | 0.024 |
| O | 26.739 |

Several adjustments were made to the databases used in the calculations to account for kinetic and other observations. Due the relatively low reaction temperatures, methane was removed from the initial list of species that could form in the Strytan system (Wang et al., 2018). Similarly, antigorite formation was suppressed since it would not occur on the short timescale and temperature range considered in this study (Wenner et al., 1973; McDermott, 2015). To match the observed inorganic carbon, Arnorsson and Andresdottir (1995) rainwater values (Supplemental Table 1), HCO_3_^-^ was adjusted until the speciation of CO_2_ in the EQ3 output file matched the measured value (total inorganic carbon of 0.00023 mol/kg). In order to validate the results of the modeled Strytan hydrothermal fluid, energetics calculations were also performed using previously measured concentrations of hydrothermal fluid from Strytan (denoted as “field sample”) (Price et al., 2017).

#### 2.2.2 Modeling water-rock reactions with EQ6

The speciated fluids, taken from the EQ3 output file, were titrated into increasing amounts of basaltic rock in the reaction path program EQ6. For the host rock, the average of six basalt samples from Iceland was used: NF-8 to NF-13 (Arnorsson et al., 2002) (Supplemental Table 1). These olivine-basalt samples (∼1% olivine average) were taken from Siglufjord (Siglufjörður in Icelandic), which is a small fjord located near the mouth of Eyjafjord, and therefore should have the same rock type (Arnorsson et al., 2002). The weight percent of each major oxide was converted into mol/kg by multiplying the average weight percent by the stoichiometric coefficient of the cation and dividing by the molecular weight of the oxide (Table 1). The molality of iron in Fe_2_O_3_ and FeO were summed to obtain total iron. The EQ6 file was run from 2 ºC to 142 ºC and the pressure was kept at 78 bar.

### 2.3 Energetics Calculations

Values of Gibbs energies of reactions (Δ*G*_*r*_) were calculated using:

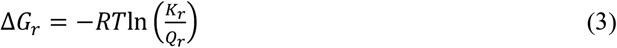

where *R* is the gas constant, *T* is temperature (Kelvin), and *K*_*r*_ represents the equilibrium constant for the reaction of interest. Values of *K*_*r*_ were calculated using the revised-HKF equations of state (Helgeson *et al*. 1981; Shock *et al*. 1992; Tanger and Helgeson 1988), the SUPCRT92 software package (Johnson *et al*. 1992), and thermodynamic data taken from a number of sources (Bricker 1965; Chase 1998; Helgeson *et al*. 1978; Hem *et al*. 1982; LaRowe and Amend 2014; Robie and Bethke 1963; Robie and Hemingway 1985; Schulte *et al*. 2001; Senoh *et al*. 1998; Shock and Helgeson 1988; Shock *et al*. 1997; Snow *et a l*. 2013). Values of *Q*_*r*_ are calculated using

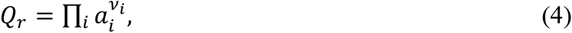

where *a*_*i*_ stands for the activity of the *i*th species and *v*_*i*_ corresponds to the stoichiometric coefficient of the *i*th species in the reaction of interest. Negative values of Δ*G*_*r*_ are said to be exergonic and positive values are endergonic; Δ*G*_*r*_ = 0 defines equilibrium. Because standard states in thermodynamics specify a composition and state of aggregation (Amend and LaRowe 2019; LaRowe and Amend 2020) values of *Q*_*r*_ must be calculated to take into account how environmental conditions impact Gibbs energy calculations. In this study, we use the classical chemical-thermodynamic standard state in which the activities of pure liquids and solids are taken to be 1 as are those for aqueous species in a hypothetical 1 molal solution referenced to infinite dilution at any temperature or pressure.

Activities are related to concentration, *C*, by

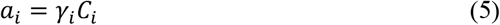

where *γ*_*i*_ and *C*_*i*_ stand for the individual activity coefficient and concentration of the *i*th species, respectively. Values of *γ*_*i*_ are computed using an extended version of the Debye-Hückel equation (Helgeson 1969).

In order to better compare the potential energy of different reactions, values of Δ*G*_*r*_ were normalized to energy per kg of H_2_O or vent fluid using the following equation:

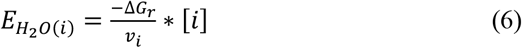

where [*i*] is the molal concentration in a kg of seawater of the limiting electron acceptor or donor. Values of 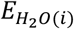 v are calculated on a log scale to facilitate order-of-magnitude differences. The normalization step is important as reactions involving reactants that have low concentrations in the fluid may have very exergonic values of Δ*G*_*r*_, but low 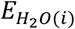 due to [*i*] being small (Price et al., 2015).

The activities of reactants and products required to calculate values of Δ*G*_*r*_ for the 85 redox reactions considered in this study (see Supplemental Table 2) were determined by reacting, in silico, rainwater with local rocks using the EQ3/6 software package. In the case of Eridania, the rainwater composition was computed based on an assumed atmospheric composition as detailed below. The water-rock calculations were conducted using three different mixing ratios of hydrothermal fluid (HF) and seawater (SW): 10 % HF: 90 % SW, 50:50, and 90 % HF: 10 % SW. For Eridania, a hypothetical Noachian basin surface water was used in place of seawater (Catling, 1999) (Supplemental Table 3).

## 3. Results

### 3.1 Strytan

The energy per kilogram of H_2_O of each reaction for SHF field data for 10:90, 50:50, and 90:10 mixing ratios are shown in Figures 1A through 1F, with different colors for each electron donor (ED) and acceptor (EA). The results are grouped by ED and EA even if the product species differ (e.g. reduced nitrogen species NH_4_^+^ and N_2_). The five most exergonic reactions occurring deep in the SHF system (90% HF:10% SW) are:

**Figure 1.**
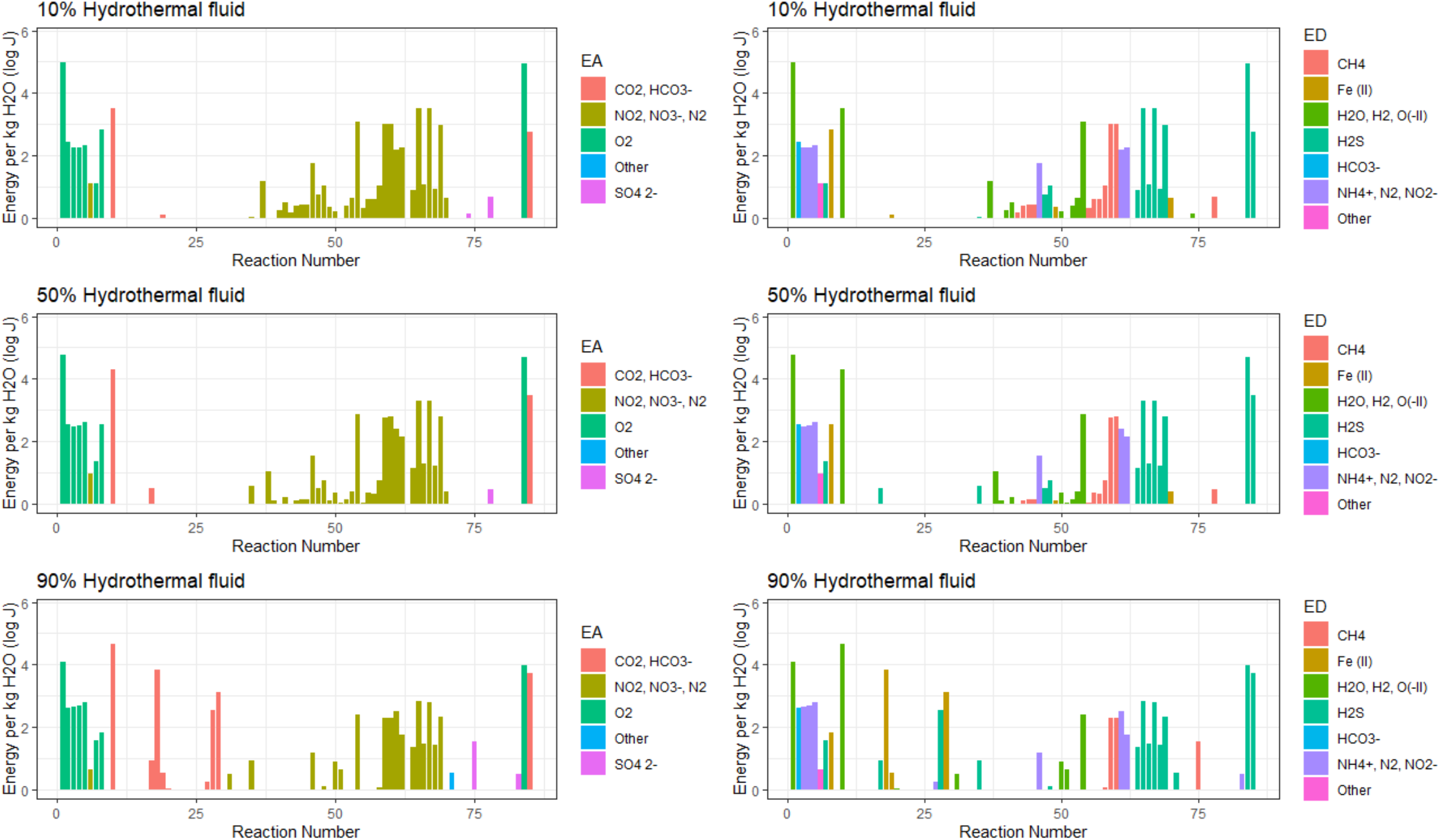
Gibbs energies of the 85 reactions considered in this study for the noted mixing ratios of hydrothermal vent fluid and seawater in the Strytan hydrothermal system. The measured chemical composition of Strytan vents was used as the hydrothermal fluid. Reactions are colored by electron acceptor or donor (left versus right side). The electron donors and acceptors are grouped when applicable with other donors or acceptors that have similar reaction equations that may differ slightly in product stoichiometry (e.g., CO_2_^-^ and HCO_3_^-^). Reactions listed as “other” represent dissociation reactions (e.g. reaction 51: 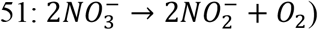).

1. Hydrogenotrophic methanogenesis (reaction 10): CO_2_ + 4H_2_ → CH_4_ + 2H_2_O
2. Aerobic hydrogen oxidation (reaction 1): 2H_2_ + O_2_ → 2H_2_O
3. Hydrogen sulfide oxidation (reaction 84): 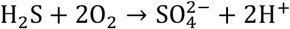
4. Pyrite oxidation coupled to carbon dioxide reduction (reaction 18):

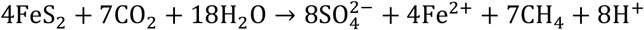
5. Hydrogen sulfide oxidation coupled to carbon dioxide reduction (reaction 85):

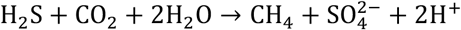

The most energetically favorable reactions in all mixing schemes for Strytan includes CO_2_ or O_2_ reduction with H_2_, or O_2_ reduction with H_2_S oxidation (Reactions 1, 10, and 84, Table 4, Supp. Table 2). These reactions are capable of producing 3-100 kJ of energy per kg of fluid in all mixing schemes (Table 4). Numerous NO_3_^-^ reduction reactions are also favorable, and NO_3_^-^ reduction with H_2_S oxidation is among the top five most energetically yielding reactions.

**Table 2.** Noachian atmospheric composition and concentration of equilibrated aqueous species in Martian rainwater.

|  | Percentage in atmosphere (%) | Partial Pressure (bar) | Concentration in rainwater (M) |
| --- | --- | --- | --- |
| **CO<sub>2</sub><sup>a</sup>** | 92.00 | 1.5 | 1.0 x 10<sup>-1</sup> |
| **H<sub>2</sub><sup>a</sup>** | 3.50 | 0.057 | 5.1 x 10<sup>-5</sup> |
| **CH<sub>4</sub><sup>a</sup>** | 3.50 | 0.057 | 1.3 x 10<sup>-4</sup> |
| **N<sub>2</sub>** | 0.25 | 0.004 | 3.8 x 10<sup>-6</sup> |
| **CO<sup>b</sup>** | 0.007 | 0.001 | 1.6 x 10<sup>-6</sup> |
| **NO** | 0.10 | 0.001 | 5.2 x 10<sup>-6</sup> |
| **SO<sub>2</sub>** | >0.007 | 0.001 | 4.7 x 10<sup>-9</sup> |
| **HCl** | >0.007 | 0.001 | 5.5 x 10<sup>-9</sup> |
| **Ar<sup>c</sup>** | 0.25 | 0.004 | n/a |
| **He<sup>c</sup>** | 0.25 | 0.004 | n/a |
| **Ne<sup>c</sup>** | 0.25 | 0.004 | n/a |
| **HCO<sub>3</sub><sup>-</sup>** | - | - | 1.0 x 10<sup>-2</sup> |
| **Cl<sup>-</sup>** | - | - | 5.5 x 10<sup>-5</sup> |
| **NO<sub>3</sub><sup>-</sup>** | - | - | 0 |
| **NO<sub>2</sub><sup>-</sup>** | - | - | 0 |
| **NH<sub>4</sub><sup>+</sup>** | - | - | 9.0 x 10<sup>-1</sup> |
| **NH<sub>3</sub>** | - | - | 1.1 x 10<sup>-6</sup> |
| **SO<sub>4</sub><sup>2-</sup>** | - | - | 4.7 x 10<sup>-9</sup> |

“n/a” indicates that the gas was not included in water composition.
<sup>a</sup>(Wordsworth et al., 2017)
<sup>b</sup>(Owen et al., 1977)
<sup>c</sup> Noble gases Ar, He and Ne were not included in the rainwater model as these gases do not contribute to ionic strength and would not have an effect on the redox chemistry of the system.

**Table 3.** Oxide content of Backstay rock and of the same rock reduced to 5% olivine.

| **Olivine**<br>**(wt %)** | Forsterite<br>(wt %) | Fayalite<br>(wt %) | Total SiO<sub>2</sub><br>(wt %) | Total MgO<br>(wt %) | Total FeO<br>(wt %) | Total Fe<sub>2</sub>O<sub>3</sub><br>(wt %) | Total Fe<br>(wt %) |
| --- | --- | --- | --- | --- | --- | --- | --- |
| **16<sup>a</sup>** | 8.7 | 7.3 | 49.6 | 8.32 | 10.7 | 3.33 | 14.03 |
| **5** | 2.7 | 2.3 | 45.6 | 4.88 | 8.04 | 2.45 | 10.49 |

<sup>a</sup>From McSween et al., 2006

**Table 4.** Most favorable reactions for each mixing ratio for the Strytan field data and Eridania thermodynamic models with 5% and 16% olivine (ol).

| SHF 10% HF + 90% SW |  |  |  | Eridania 10% HF + 90% BW (5% Ol) |  |  |  | Eridania 10% HF + 90% BW (16% Ol) |  |  |  |
| --- | --- | --- | --- | --- | --- | --- | --- | --- | --- | --- | --- |
| Reaction | EA | ED | Log J | Reaction | EA | ED | Log J | Reaction | EA | ED | Log J |
| 1 | O<sub>2</sub> | H<sub>2</sub> | 4.99 | 12 | CO<sub>2</sub> | NH<sub>4</sub><sup>+</sup> | 6.74 | 12 | CO<sub>2</sub> | NH<sub>4</sub><sup>+</sup> | 6.74 |
| 85 | O<sub>2</sub> | H<sub>2</sub>S | 4.94 | 11 | CO<sub>2</sub> | NH<sub>4</sub><sup>+</sup> | 5.83 | 11 | CO<sub>2</sub> | NH<sub>4</sub><sup>+</sup> | 5.83 |
| 66 | NO<sub>3</sub><sup>-</sup> | H<sub>2</sub>S | 3.51 | 18 | CO<sub>2</sub> | Fe (II) | 5.81 | 18 | CO<sub>2</sub> | Fe (II) | 5.82 |
| 68 | NO<sub>3</sub><sup>-</sup> | H<sub>2</sub>S | 3.51 | 23 | HCO<sub>3</sub><sup>-</sup> | NH<sub>4</sub><sup>+</sup> | 5.38 | 23 | HCO<sub>3</sub><sup>-</sup> | NH<sub>4</sub><sup>+</sup> | 5.38 |
| 10 | CO<sub>2</sub> | H<sub>2</sub> | 3.50 | 86 | CO<sub>2</sub> | H<sub>2</sub>S | 4.82 | 86 | CO<sub>2</sub> | H<sub>2</sub>S | 4.82 |
| SHF 50% HF + 50% SW |  |  |  | Eridania 50% HF + 50% BW (5% Ol) |  |  |  | Eridania 50% HF + 50% BW (16% Ol) |  |  |  |
| Reaction | EA | ED | Log J | Reaction | EA | ED | Log J | Reaction | EA | ED | Log J |
| 1 | O<sub>2</sub> | H<sub>2</sub> | 4.75 | 12 | CO<sub>2</sub> | NH<sub>4</sub><sup>+</sup> | 6.34 | 12 | CO<sub>2</sub> | NH<sub>4</sub><sup>+</sup> | 6.35 |
| 85 | O<sub>2</sub> | H<sub>2</sub>S | 4.68 | 14 | CO<sub>2</sub> | N<sub>2</sub> | 6.31 | 14 | CO<sub>2</sub> | N<sub>2</sub> | 6.29 |
| 10 | CO<sub>2</sub> | H<sub>2</sub> | 4.30 | 18 | CO<sub>2</sub> | Fe (II) | 5.92 | 18 | CO<sub>2</sub> | Fe (II) | 5.89 |
| 86 | CO<sub>2</sub> | H<sub>2</sub>S | 3.46 | 11 | CO<sub>2</sub> | NH<sub>4</sub><sup>+</sup> | 5.37 | 11 | CO<sub>2</sub> | NH<sub>4</sub><sup>+</sup> | 5.44 |
| 66 | NO<sub>3</sub><sup>-</sup> | H<sub>2</sub>S | 3.30 | 25 | HCO<sub>3</sub><sup>-</sup> | N<sub>2</sub> | 5.26 | 25 | HCO<sub>3</sub><sup>-</sup> | N<sub>2</sub> | 5.26 |
| SHF 90% HF + 10% SW |  |  |  | Eridania 90% HF + 10% BW (5% Ol) |  |  |  | Eridania 90% HF + 10% BW (16% Ol) |  |  |  |
| Reaction | EA | ED | Log J | Reaction | EA | ED | Log J | Reaction | EA | ED | Log J |
| 10 | CO<sub>2</sub> | H<sub>2</sub> | 4.67 | 18 | CO<sub>2</sub> | Fe (II) | 5.80 | 14 | CO<sub>2</sub> | N<sub>2</sub> | 5.86 |
| 1 | O<sub>2</sub> | H<sub>2</sub> | 4.07 | 14 | CO<sub>2</sub> | N<sub>2</sub> | 5.79 | 18 | CO<sub>2</sub> | Fe (II) | 5.76 |
| 85 | O<sub>2</sub> | H<sub>2</sub>S | 3.99 | 12 | CO<sub>2</sub> | NH<sub>4</sub><sup>+</sup> | 5.70 | 12 | CO<sub>2</sub> | NH<sub>4</sub><sup>+</sup> | 5.70 |
| 18 | CO<sub>2</sub> | Fe (II) | 3.82 | 11 | CO<sub>2</sub> | NH<sub>4</sub><sup>+</sup> | 5.59 | 11 | CO<sub>2</sub> | NH<sub>4</sub><sup>+</sup> | 5.46 |
| 86 | CO<sub>2</sub> | H<sub>2</sub>S | 3.73 | 29 | HCO<sub>3</sub><sup>-</sup> | Fe (II) | 4.12 | 25 | HCO<sub>3</sub><sup>-</sup> | N<sub>2</sub> | 4.50 |

Furthermore, the maximum amount of energy produced by the most favorable reaction consistently decreases as the amount of hydrothermal fluid increases in the mixing ratio. However, the amount of energy an individual reaction can produce varies between mixing ratios (e.g., an increase in energy with CO_2_ reduction with H_2_ as the percentage of HF increases) (Table 4). Generally, reductions with O_2_ as the electron acceptor are favorable at all mixing ratios (shown in green; Figure 1). Reduction of nitrogen-bearing species are overall mostly favorable at 10 and 50% HF and decrease in potential energy at 90% HF (shown in yellow; Figure 1). An inverse relationship is seen with CO_2_ reduction reactions, where more reactions become favorable at 90% HF in the field sample (shown in red; Figure 1). Only two reactions for the reduction of SO_4_^2-^ are favorable at any given mixing ratio, with Reaction 75, the reduction of SO_4_^2-^ with CH_4_, being the only reaction to produce more than 10 joules of energy per kg fluid in the 90% HF calculation.

### 3.2. Recreating Strytan vent fluid chemistry from in silico

To test the reliability of EQ3/6 to determine the chemical composition of fluids on Noachian Mars resulting from water-rock interactions, we calculated the composition of Strytan hydrothermal fluids based on the composition of the rainwater and the rocks that it reacted with and compared them to actual measured compositions of the vent fluid. The measured and modeled compositions of the end-member hydrothermal fluids for the SHF are shown in Supplemental Table 4. The final modeled fluid did not contain O_2_ or nitrogen-bearing species. The concentrations of H_2_, H_2_S, SO_4_^2-^ species in the model fluid and field data values are on the same order of magnitude, except for Fe^2+^, which was much lower in the modeled fluids (1.48 × 10^−7^ and 3.22 × 10^−12^ mol/kg, respectively). The modeled hydrothermal fluid had a pH of 9.79, which closely matches the actual SHF pH of 10.

As an additional step for testing the reliability of our modeling approach, we also calculated values of Δ*G*_*r*_ for the same set of reactions in SHF modeled fluids as those for measured vent data. The results are shown in Supplemental Figure 1. This ground-truthing experiment shows that the modeled and field-based Gibbs energy calculations are similar, with the 10:90 mixing ratio results most closely matching the field data energetics (Supplemental Fig. 1). The model results begin to diverge more from the field data at the 50%, and even more in the 90%, HF mixing ratio. Within the 90% HF model, the largest energetics differences involved the electron acceptors O_2_, CO_2_ / HCO_3_^-^, and NO_2_^-^ / NO_3_^-^ / N_2_, and the electron donors NH_4_^+^ and Fe^2+^ (and one H_2_ oxidation reaction; Supplemental Figure 1). Some NO_3_^-^ reduction reactions, coupled to H_2_S oxidation, were also different by about 10 joules. The overall potential energy of the 90% HF model is, for the most part, lower than the 90% HF field sample (Supplemental Figure 1).

However, the most favorable reactions at all mixing ratios for the model are similar to the field data, with CO_2_ or O_2_ reduction coupled with H_2_ or H_2_S oxidation being the most favorable (reactions 1, 10 and 85). Overall, the SHF model reproduces the results from the calculations using sampled Strytan fluid, providing confidence that similar water-rock modeling can be reliably applied to study the potential energetics of ancient vent system at Eridania.

### 3.3. Eridania

The energy per kilogram of H_2_O of each reaction for the Eridania model with 5% or 16% olivine content are shown in Figures 2-3 for the 10:90, 50:50, and 90:10 mixing ratios, colored by electron donor (ED) and acceptor (EA). The end-member hydrothermal fluid composition of both the 5% and 16% olivine models can be found in Supplemental Table 4. The most energetically favorable reactions involve almost exclusively CO_2_ and HCO_3_^-^ as electron acceptors and NH_4_^+^, N_2_ and H_2_ as electron donors (Reactions 10-12, 14, 21, 23, 25 28-29, 31-34, 36-37, and 85). There were no significant differences between the Gibbs energy yields for the two rock types considered at 10% HF:90% SW fluid in Eridania, likely due to the small proportion of hydrothermal fluid included in this calculation. At 90% HF, however, the Gibbs energies for both rock models became more variable, with approximately 60% of the favorable reactions having more energy in the 5% olivine model (9 out of 16) (Figures 2 and 3). These energetic differences range from small, less than 5% differences, to more than 1000 joules. One reason for the variability in energy for the two olivine models is the difference in mineralogy resulting in subtle compositional differences in the hydrothermal fluid output, such as an increase in Fe_2_^+^ in the 5% olivine model and NH_4_^+^ in the 16% olivine model (Supplemental Table 2).

**Figure 2.**
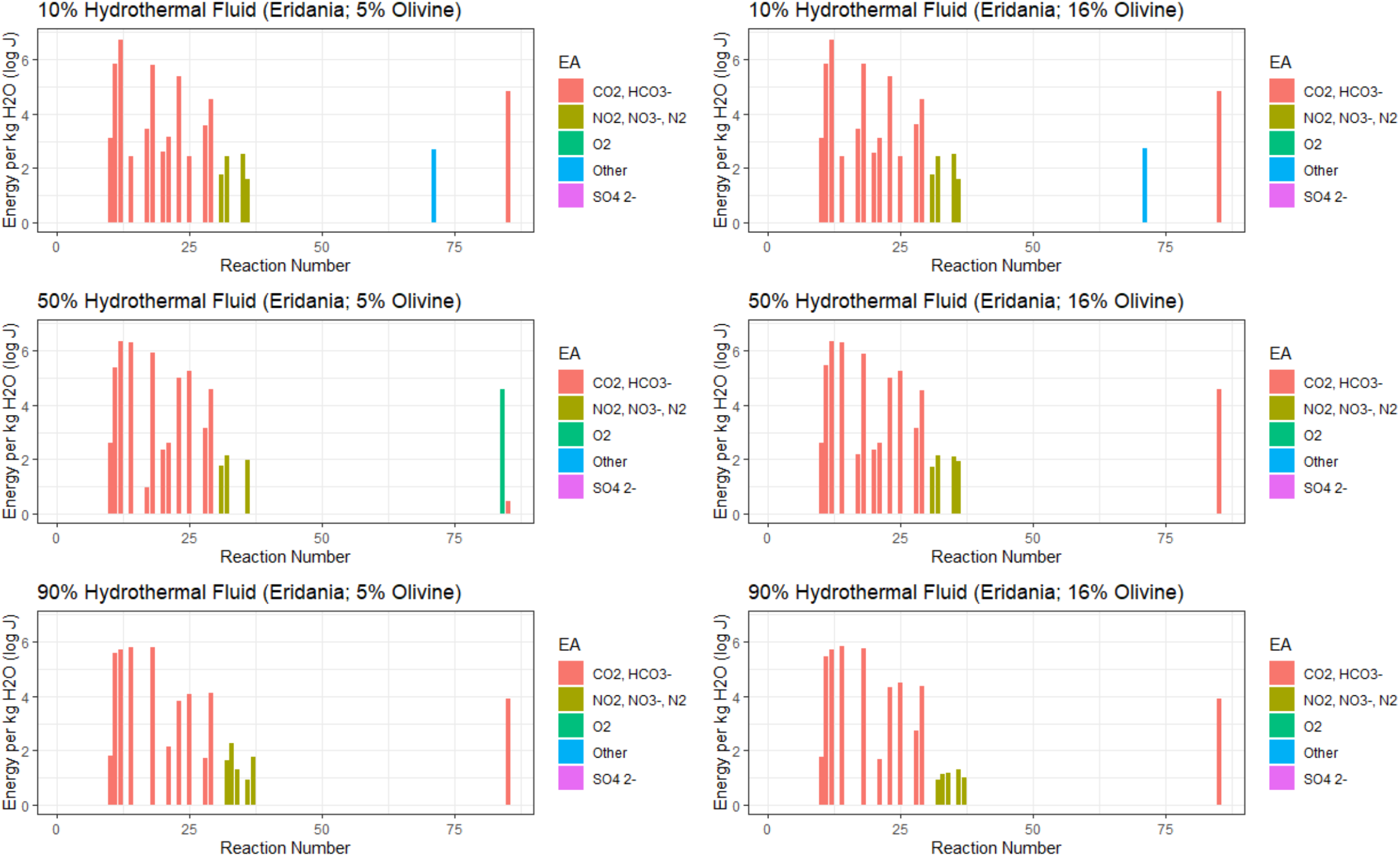
Gibbs energies of the 85 reactions considered in this study for the Eridania models containing 5% or 16% olivine. Reactions are colored by electron acceptor.

**Figure 3.**
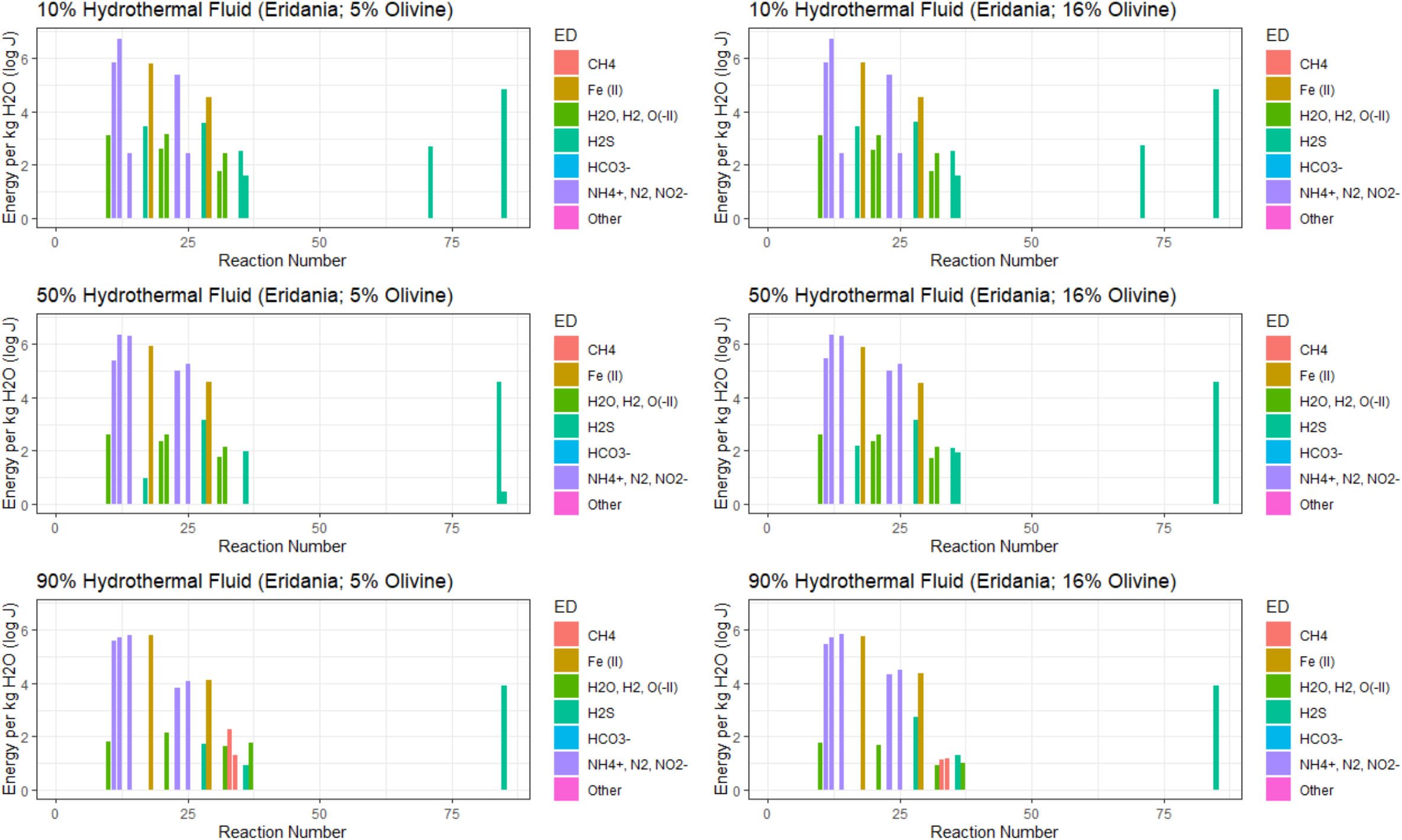
Gibbs energies of the 85 reactions considered in this study for the Eridania models containing 5% or 16% olivine. Reactions are colored by electron acceptor.

For both olivine models, the amount of energy a favorable reaction can yield decreases as the percentage of hydrothermal fluid increases (Table 4). While the potential energy values differ between the 5% and 16% olivine models, the types of favorable reaction between the models remains the same. The favorable reactions at Eridania consist mainly of anaerobic processes such as CO_2_/HCO_3_^-^ reduction reactions, with N_2_ reduction producing less energy on average (Table 4). Favorable electron donors include NH_4_^+^, H_2_S, Fe^2+^ and H_2_. For all mixing ratios, no reaction involving SO_4_^2-^ was favorable. Reactions involving the reduction of NO_3_^-^ or NO_2_^-^, were highly unfavorable at all mixing ratios for the Eridania model, as these reactants are limited by the slightly reducing composition of the atmospheric model. Thermodynamic speciation calculations determined that NO_3_^-^ and NO_2_^-^ would be essentially absent from Martian rainwater at pH 4 due to the low abundance of NO in the atmospheric model (0.10%). The speciation calculations instead favored the presence of more reduced forms of nitrogen, such as NH_4_^+^ and N_2_, at the chosen pH and log *f*O_2_ values. It is important to note that many of the O_2_ reduction reactions at Eridania had favorable Δ*G*_*r*_ values (i.e., negative values), but failed to produce even trace amounts of energy per kg H_2_O values due to the limiting reactant step in this calculation (equation 7). Since O_2_ is extremely low in the atmospheric model, its low abundance limits its potential as an electron acceptor when put into an environmental context. These results further support the use of the limiting reactant normalization step in energetics studies (Lu et al., 2020).

For putative microorganisms living closest to the site of active venting, the 90% HF mixing ratio scenario, the top five exergonic reactions for both the 5% and 16% olivine models are:

1. Methanogenesis coupled to dinitrogen oxidation (nitrite production) (reaction 14):

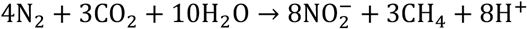
2. Pyrite oxidation coupled to CO_2_ reduction (reaction 18):

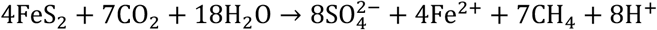
3. Ammonium-oxidation dependent Methanogenesis (nitrite production) (reaction 12):

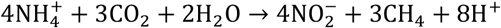
4. Ammonium-oxidation dependent Methanogenesis (dinitrogen production) (reaction 11):

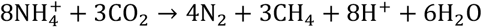
5. Methanogenesis coupled to dinitrogen oxidation and bicarbonate reduction (reaction 25):

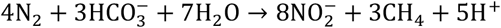

The Eridania model also produced less favorable reactions as compared to SHF, with the 90% HF models for Eridania having 16 favorable reactions, as compared to Strytan’s 39 (Figure 4). Reactions in the Eridania model typically produce a higher energy yield than the same reaction at SHF (Figure 4).

**Figure 4.**
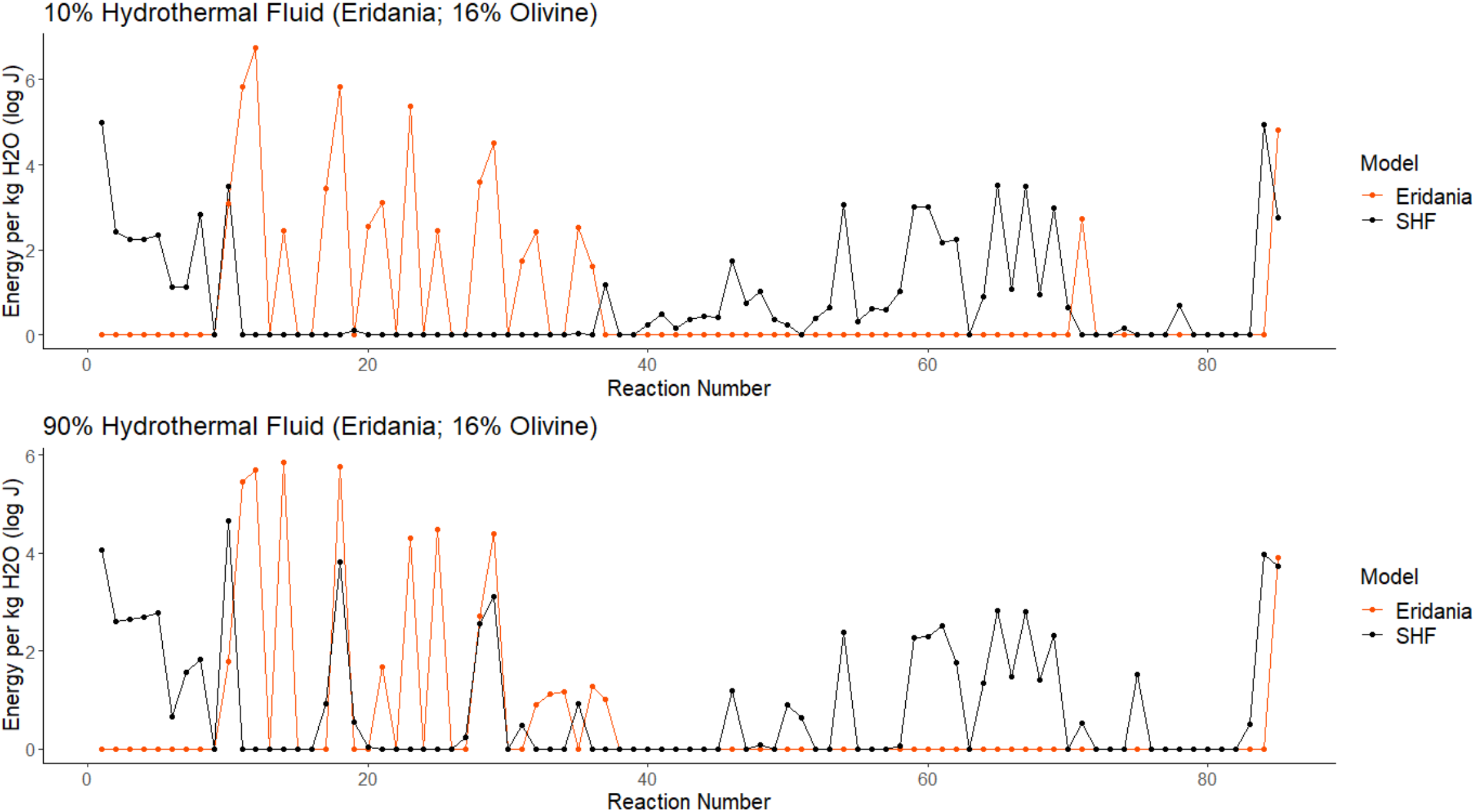
Overlay of the Gibbs energies for reactions able to produce more than 1 joule of energy (i.e., favorable) for the Strytan and Eridania models at 10% and 90% hydrothermal fluid mixing ratios. For Eridania, the 16% olivine model was used.

## 4. Discussion

Our results show that if microorganisms existed in Eridania Basin on Noachian Mars, they could have been supported by methane-forming reactions (*e.g*. reactions 11, 12, 14, 18, and 25) to the tune of 15-700 kilojoules per kilogram of fluid for the most favorable reactions.

Reaction 18, pyrite oxidation coupled to carbon dioxide reduction, is highly favorable for both Eridania and in the SHF analogue site in Northern Iceland, producing two orders of magnitude more energy in the Eridania model than SHF (600 and 6 kJ, respectively). In order to compare Eridania and SHF as a potential analogue, a closer analysis into the energy potentials of each site is considered below.

### 4.1. The energetic potential of the Strytan Hydrothermal Field

Our results suggest that the SHF can theoretically support a diversity of metabolisms. In particular, hydrogenotrophy may play a crucial role, as it is energetically favorable in the 50% and 90% HF mixing ratios, followed by the aerobic oxidation of hydrogen or hydrogen sulfide (reactions 2 and 3). Of the predicted reactions, the catabolic pathways represented by reactions 10, 1 and 85 have been extensively described and studied in a number of different *Archaea* (reaction 10) as well as *Bacteria* (reactions 1 and 84), and are expected to occur within the SHF.

Results from previous papers on the microbial communities of Big Strytan support the role of H_2_ oxidation reactions in all mixing schemes in our study with the detection of abundant members of the *Aquificaee* phylum (Marteinsson et al., 2001; Twing et al., in submission). *Aquificae* are thermophilic bacteria that can be found in many hydrothermal systems, such as shallow and deep-sea vents as well as terrestrial hot springs (Ferrera et al., 2007; Giovannelli et al., 2017). All cultured members of the *Aquificae* are chemolithoautotrophs capable of hydrogen or hydrogen sulfide oxidation coupled either to oxygen (microaerophilic in the *Aquificaceae* family), nitrate or elemental sulfur reduction (in the other two families of the phylum) (Bonch-Osmolovskaya, 2008; Giovannelli et al., 2017). The minimum and maximum growth temperature for most *Aquificales* ranges between 60-95 ºC, in the range to the 50% and 90% hydrothermal mixing temperatures at SHF (40 and 66 ºC, respectively) (Bonch-Osmolovskaya, 2008). It is likely that two genera of *Aquificaceae, Thermocrinis* (Twing et al., in submission and *Hydrogenobacter* (Marteinsson et al., 2001) dominate Strytan hydrothermal chimneys. Members of the genus *Hydrogenobacter* are common in both terrestrial hot springs and shallow-water hydrothermal vents and have an energy metabolism comparable to the genus *Thermocrinis* (Price and Giovannelli, 2017). It is possible that members of the *Aquificae* form a biofilm on the vent precipitates and obtain energy from the mixing of the vent fluids with seawater (i.e., similar to the 50-90% HF mixing ratios) taking advantage of the availability of hydrogen and hydrogen sulfide as electron donors in the vent fluids while using oxygen from seawater.

While the three most exergonic reactions noted above are reasonable based on known microbial physiology, reactions 4 and 5 have never been reported for known groups of *Bacteria* and *Archaea*. To date, pyrite oxidation and sulfide oxidation have been reported coupled to either oxygen or nitrate reduction, and the only methane producing reactions known are performed by members of the *Euryarchaeota* phylum using either hydrogen, methanol or acetate as electron donors (Berghuis et al., 2019), and only recently has the aerobic production of methane has been proposed for a *Bacteria* (Wang et al., 2021). Additionally, the reduction of CO_2_ as an electron acceptor is currently known only in methanogens and acetogens. It is therefore unlikely that the predicted reactions generating methane (4 and 5, above) from the pyrite- or sulfide-dependent reduction of CO_2_ will be confirmed as being present in SFH in future microbiological studies. Despite this, metabolic reactions predicted to be theoretically favorable have been discovered in extant microbes in the past (*e.g*., the anammox reaction, (Kuenen, 2008); anaerobic oxidation of methane (reviewed in Knittel and Boetius, 2009)) and are actively being searched today (Amend et al. 2020; LaRowe et al., 2021). It is thus possible that similar metabolism might be present in uncultured microbes in SHF, and in-depth thermodynamic-informed microbial investigations might elucidate the role of these metabolisms in the future.

### 4.2 The energetic potential of the Noachian Eridania hydrothermal field

Within the favorable reaction list for Eridania, methane-producing reactions are prevalent. These results suggest that potential microbes living around an Eridania hydrothermal vent system would likely have been methanogens utilizing electron donors such as NH_4_^+^, FeS_2_, and H_2_. The likelihood of these catabolisms can only be evaluated in context of Earth microbiology. As discussed above, currently only three types of methanogenesis are known to occur on extant Earth (hydrogenotrophic, methylotrophic and acetoclastic). To our knowledge, N_2_ oxidation, iron sulfide oxidation and ammonia oxidation have never been reported in conjunction with methane formation, and never with CO_2_ or HCO_3_^-^ as electron acceptor. As mentioned above, the reduction of CO_2_ as electron acceptor is currently known only in methanogens and acetogens. Additionally, N_2_ has never been reported as electron donor or terminal electron acceptor. Despite this, our current knowledge of the diversity of redox couples used by biology is limited to extant biology (Jelen et al., 2016; Moore at al., 2017). It is possible that some of the predicted reaction might have been used by life in the past (Moore at al., 2017), or might be discovered within the large fraction of uncultured microbes in the future (Lloyd et al., 2018).

### 4.3 Strytan as an analogue for a Noachian Eridania vent system

The Strytan hydrothermal system is fed by rainwater that percolates through an anomalously heated basaltic aquifer. Due to water-rock reactions, this aquifer becomes anoxic and alkaline, with high concentrations of dissolved SiO_2_, and measurable amounts of H_2_ and CH_4_. The putative hydrothermal system in ancient Eridania is thought to have also had rainwater as a source fluid (i.e., both would have been fresh) and be anoxic and warm, likely with high SiO_2_ due to reaction with basalt and saponite precipitates. The major difference between these systems is O_2_.

From a catabolic standpoint, the main similarity between Strytan and Eridania is the favorable energetics of CO_2_/HCO_3-_ reduction reactions in the 90% HF mixing ratios (reactions 10, 18, 28, 29, 85). These reactions use H_2_S, FeS_2_ and H_2_ as electron donors (Supplemental Table 1). All of these reactions produce methane and therefore suggest that Eridania supported methanogens. However, while methanogenesis is the main metabolism expected at Eridania, SHF is able to support a more diverse list of potential metabolisms (Figure 4).

Both Strytan and Eridania exhibit an overall decrease in Gibbs energies for a given reaction from the 10% to 90% HF mixing ratio. This decrease in potential energy as hydrothermal fluid content increases is due to the seawater and basin water compositions being more chemically diverse than the hydrothermal fluids. For example, at SHF, concentrations of O_2_, HCO_3_^-^ and N_2_ can be used for seawater calculations, whereas these same compounds were below detection for the hydrothermal fluid. When the 90% SW calculation is performed, a compound found exclusively in seawater, such as O_2_, will have a higher concentration than in the 10% SW calculation. The higher SW% calculations (i.e., lower HF%) include more chemical species measured only in seawater, such as O_2_, and can therefore produce more energy. The basin water model for Mars is in direct contact with the atmosphere and therefore contains significantly higher concentrations of CO_2_, HCO_3_^-^, H_2_S, and N_2_ and thus can support higher energy yields at a higher proportion of basin water in the mixing ratio.

Our results indicate that modern Strytan should only be considered as an analogue for Eridania for methane producing reactions such as 10, 18, 28, 29, 85 at 90% HF. None of these reactions involve O_2_. Additionally, the most favorable reaction at SHF, methanogenesis (reaction 10), would produce 93% less energy than the most favorable reaction in both Eridania models (reactions 14 and 18), with greater than an order of magnitude difference between the two sites. Overall, the most favorable reactions at Eridania have higher energy potentials than SHF by an order of magnitude (630 kJ per kg of fluid vs 46 kJ per kg of fluid), most likely because the reactions that are favorable at Eridania utilize reactants that are much higher in concentration than at Strytan. For example, one of the most favorable ED at Eridania, NH_4_^+^ was calculated to be 0.575 mol/kg for the Eridania models, whereas NH_4_^+^ was only measured to be 1.55×10^−6^ mol/kg at Strytan. Eridania and Strytan also differ in the number of favorable reactions, with SHF having double the number of favorable reactions at the 90% HF mixing ratio (Figure 4).

The reduced number of favorable reactions and lack of diversity in the types of favorable electron acceptors and donors compared to Strytan are a result of the low concentrations of O_2_, NO_3_^-^ and NO_2_^-^ in the atmospheric model, which limit their ability to act as electron acceptors (Figures 2-3, Supplemental Figure 1). Another difference between SHF and Eridania is the end-member pH of the hydrothermal fluid, with Eridania’s pH being approximately 2 pH units lower (7.96 in the 5% olivine model vs 10.03 in the field). The difference is likely due to the buffering effect seen at SHF, with CaCO_3_ precipitation exhausting the CO_2_ groundwater buffer and allowing the fluids to reach a higher pH (Price et al., 2017). This would not occur at Eridania, where CO_2_ is more dominant in the atmosphere and the meteoric source fluid.

Previous studies on the energetics of ancient Mars have focused only on a small number of reactions (e.g., Shock, 1997; Varnes et al., 2003; Marlow et al., 2014). Varnes et al. (2003) used energetics calculations to determine the energy yield for two reactions, methanogenesis and sulfate reduction, and concluded that there would be sufficient energy available for these metabolisms. However, they used a host rock derived from Martian meteorites, which have a nondescript origin. Similarly, Shock (1997) and Marlow et al. (2014) investigated the energetics of methanogenesis and anaerobic methane oxidation coupled with sulfate reduction (AOM), respectively, on ancient Mars. These studies also concluded that both methanogenesis and AOM would be exergonic in an ancient Mars hydrothermal system or subsurface environment.

Our results support the hypothesis that methanogenesis, using NH_4_^+^ or N_2_, would have been a potential metabolism at Eridania, and H_2_S oxidation when CO_2_ is used as the oxidant. It is important to note that Varnes et al. (2003) produced similar energy yields for methanogenesis using both a low log *f*_O2_ value similar to this study (−74 vs -80, respectively) and a high log *f*_O2_ value (−5). Furthermore, AOM and sulfate reduction failed to produce energy in the Eridania model due to the low abundance of SO_4_^2-^ in the model hydrothermal fluid. Additionally, the Backstay rock sample used did not contain any sulfur species, whereas elemental sulfur and SO_3_^-^ were measured in the meteorite samples used in Varnes et al. (2003) and likely resulted in increased favorability of sulfate reduction in their model. Any differences in reaction favorability between these studies and the current study are not necessarily surprising, as the composition of the host rock in Earth-based hydrothermal vent systems, as well as the pH and temperature at which water-rock reactions occur, can lead to variation in energetic profiles between vent systems (Amend et al., 2011).

## 5. Conclusions and Future Directions

This study is the first to provide an in-depth evaluation of the potential energy available from water-rock reactions using Noachian aged olivine-rich basalts on Mars. The knowledge gained from this study provides insights into the habitability of ancient Mars and the energetics of saponite-precipitating hydrothermal systems.

By using both published field data and thermodynamic modeling, we determined the energetics for chemolithotropic metabolism at SHF and a potential hydrothermal vent system on ancient Mars. Our results indicate that saponite precipitating hydrothermal vents are capable of maintaining chemical gradients with energetically favorable reaction potentials. While Eridania supports a smaller number of metabolically relevant redox reactions compared to SHF, the reactions that are favorable are highly exergonic. Since the SHF model underestimated the potential energy as compared to the energy calculated using the field data, it is possible that the favorable reactions at Eridania would be even more exergonic than reported in this study.

Therefore, the energetics calculations support Eridania as a potentially habitable environment for microorganisms. It should, however, be kept in mind that the potentials calculated here are for chemolithotrophy only, and do not consider heterotrophy. These results suggest that if microbes did live in the Eridania basin during the Noachian, then the lack of diversity in favorable reaction types may have resulted in potentially less diverse populations of microbes, as compared to Strytan. Microorganisms within an Eridania hydrothermal vent would likely be obligate anaerobes, whereas the mixing of oxygenated seawater at Strytan allows for a wider range of potential metabolisms depending on the degree of mixing / proximity to active venting.

However, the deep subsurface of the anaerobic basaltic aquifer at Strytan may have similar characteristics compared to Eridania. Therefore, only a few of the energetically favorable reactions determined for SHF can be considered as good analogs for the Noachian Eridania Basin hydrothermal system. These metabolisms include methane production using CO_2_ or HCO_3_^-^ and H_2_, FeS_2_, or H_2_S, and while some of these metabolisms have never been observed on extant Earth life, Gibbs energy calculations suggests that they are favorable and future research might discover them in the large fraction of uncultured microbes associated with these shallow-water hydrothermal vent systems.

Future research could consider coupled reaction networks where the product from reaction A can be utilized as a reactant in reaction B. Additionally, the results from the study can be further expanded upon by utilizing the flow rate of venting fluids at Strytan and creating reactive transport models for SHF and Eridania. The reactive transport model results will provide more information about how much energy can be harvested from a reaction given the flow rate of the hydrothermal system. A bioenergetics evaluation using ultramafic rocks at Eridania would provide additional insights. Ultramafic systems hosted by rocks with a higher olivine content, such as dunites, may also support higher energetic potentials within this thermodynamic model (McSween, 2015). The energetic calculations of this study could also be expanded to include the energy available for chemoorganoheterotrophs that utilize organic carbon sources (i.e., heterotrophs).

## Acknowledgments

The authors thank J. Hurowitz, A. Fraeman, D. Rogers, and J. Michalski for answering questions related to model design for this article. We thank Erlendur Bogasonn of the Strytan Dive Center for invaluable assistance during sampling.

## Authorship confirmation statement

**Holly Rucker**: Methodology, Formal analysis, Investigation, Data Curation, Writing - Original Draft, Writing - Review & Editing, Visualization. **Tucker Ely**: Methodology, Software, Resources, Writing - Review & Editing. **Douglas LaRowe**: Conceptualization, Methodology, Software, Resources, Writing – Review & Editing. **Donato Giovannelli**: Writing – Review & Editing. **Roy Price:** Conceptualization, Supervision, Funding acquisition, Writing – Original Draft, Writing - Review & Editing

