## Supplemental for "Quantifying the bioavailable energy in an ancient hydrothermal vent on Mars and a modern Earth-based analogue"

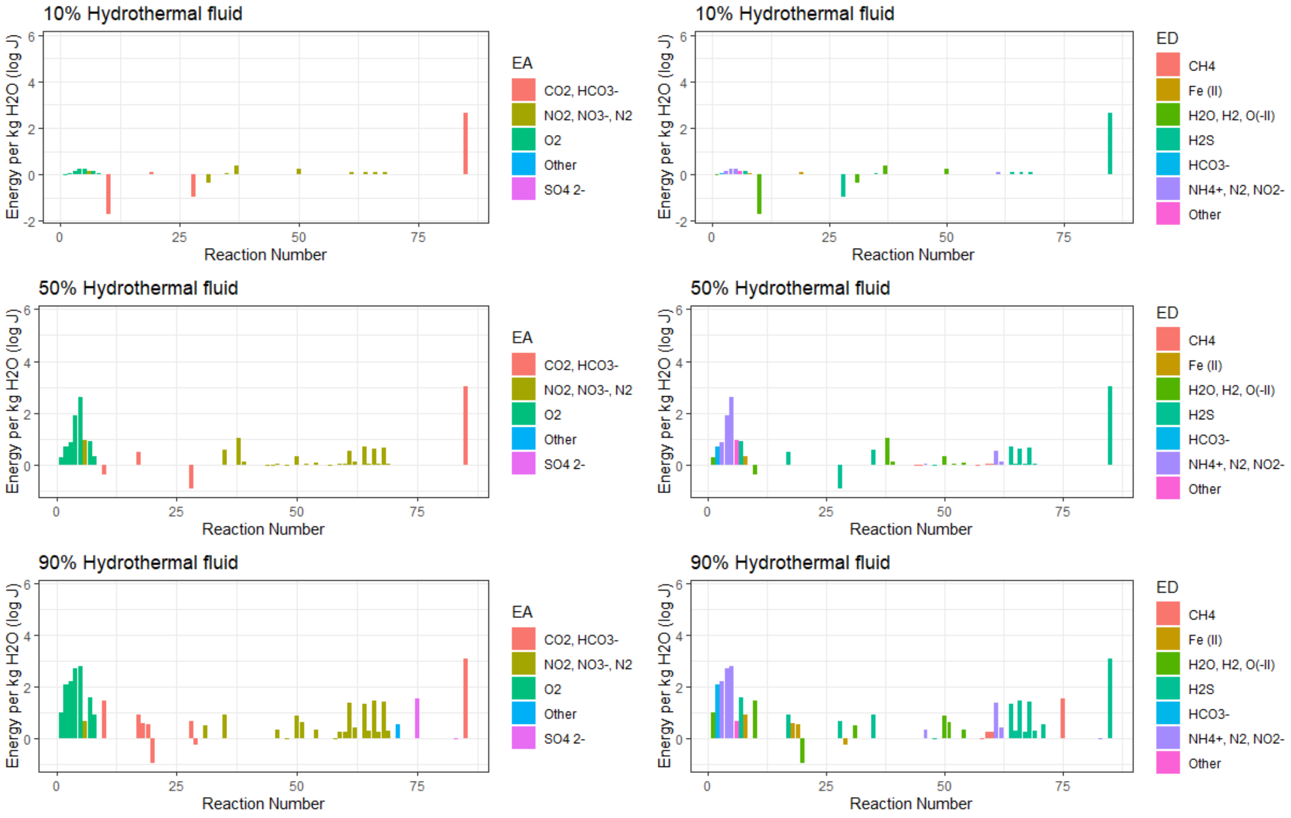


**Supplemental Figure 1**. Difference in Gibbs energies when the fluid composition of Strytan hydrothermal fluids used in the calculations was based on measured versus modeled values. The modeled energy values were subtracted from the Strytan field derived results and therefore positive energy represents more available energy in the field sample whereas negative energy indicates more energy available in the model calculations.

**Supplemental Table 1. Icelandic rainwater composition and basalltic oxide content**

| | Rainwater input[a] (mol/L) | Major Oxides | Basalt input[b] (mol/kg) |
|---|---|---|---|
| **pH** | 5.55 | **$SiO_2$** | 7.96 |
| **T ($^oC$)** | 7 | **$TiO_2$** | 0.22 |
| **$SiO_2$** | $3.16 \times 10^{-5}$ | **$Al_2O_3$** | 1.40 |
| **B** | $5.46 \times 10^{-7}$ | **$Fe_2O_3$** | 0.25 |
| **Na** | $1.87 \times 10^{-4}$ | **FeO** | 1.14 |
| **K** | $7.19 \times 10^{-5}$ | **MnO** | 0.03 |
| **Ca** | $3.37 \times 10^{-5}$ | **MgO** | 1.78 |
| **Mg** | $4.85 \times 10^{-5}$ | **CaO** | 2.03 |
| **Fe** | $1.09 \times 10^{-6}$ | **$Na_2O$** | 0.38 |
| **Al** | $1.22 \times 10^{-6}$ | **$K_2O$** | 0.02 |
| **$CO_2$** | $2.29 \times 10^{-4}$ | **$P_2O_5$** | 0.01 |
| **$SO_4^{2-}$** | $2.09 \times 10^{-5}$ | | |
| **Cl** | $2.74 \times 10^{-4}$ | | |
| **F** | $9.47 \times 10^{-7}$ | | |

[a]Values from Arnorsson and Andresdottir, 1995

[b]Values from Arnorsson et al., 2002

**Supplemental Table 2. Reaction list**

| | | |
|---|---|---|
| **Oxidation of H,C,N, and S with $O_2$** | | |
| B1 | 1 | $2H_2 + O_2 \rightarrow 2H_2O$ |
| B4 | 2 | $CH_4 + 2O_2 \rightarrow HCO_3^- + H_2O + H^+$ |
| B7 | 3 | $4NH_4^+ + 3O_2 \rightarrow 2N_2 + 4H^+ + 6H_2O$ |
| B8 | 4 | $2NH_4^+ + 3O_2 \rightarrow 2NO_2^- + 4H^+ + 2H_2O$ |
| B9 | 5 | $NH_4^+ + 2O_2 \rightarrow NO_3^- + 2H^+ + H_2O$ |
| B12 | 6 | $2NO_2^- + O_2 \rightarrow 2NO_3^-$ |
| B13 | 7 | $4H_2S + O_2 + 2Fe^{2+} \rightarrow 2FeS_2 + 4H^+ + 2H_2O$ |
| B18 | 8 | $2FeS_2 + 7O_2 + 2H_2O \rightarrow 4SO_4^{2-} + 2Fe^{2+} + 4H^+$ |
| B21 | 9 | $6Fe^{2+} + O_2 + 6H_2O \rightarrow 2Fe_3O_4 + 12H^+$ |
| **Reduction of $CO_2$** | | |
| D3 | 10 | $CO_2 + 4H_2 \rightarrow CH_4 + 2H_2O$ |
| D5 | 11 | $8NH_4^+ + 3CO_2 \rightarrow 4N_2 + 3CH_4 + 8H^+ + 6H_2O$ |
| D6 | 12 | $4NH_4^+ + 3CO_2 + 2H_2O \rightarrow 4NO_2^- + 3CH_4 + 8H^+$ |
| D7 | 13 | $NH_4^+ + CO_2 + H_2O \rightarrow NO_3^- + CH_4 + 2H^+$ |
| D11 | 14 | $4N_2 + 3CO_2 + 10H_2O \rightarrow 8NO_2^- + 3CH_4 + 8H^+$ |
| D12 | 15 | $4N_2 + 5CO_2 + 14H_2O \rightarrow 8NO_3^- + 5CH_4 + 8H^+$ |
| D15 | 16 | $4NO_2^- + CO_2 + 2H_2O \rightarrow 4NO_3^- + CH_4$ |
| D17 | 17 | $8H_2S + CO_2 + 4Fe^{2+} \rightarrow 4FeS_2 + CH_4 + 8H^+ + 2H_2O$ |
| D26 | 18 | $4FeS_2 + 7CO_2 + 18H_2O \rightarrow 8SO_4^{2-} + 4Fe^{2+} + 7CH_4 + 8H^+$ |
| D33 | 19 | $12Fe^{2+} + CO_2 + 14H_2O \rightarrow 4Fe_3O_4 + CH_4 + 24H^+$ |
| **Reduction of $HCO_3^-$** | | |
| E2 | 20 | $HCO_3^- + H_2O + H^+ \rightarrow CH_4 + 2O_2$ |
| E3 | 21 | $4H_2 + HCO_3^- + H^+ \rightarrow CH_4 + 3H_2O$ |
| E5 | 22 | $8NH_4^+ + 3HCO_3^- \rightarrow 4N_2 + 3CH_4 + 5H^+ + 9H_2O$ |
| E6 | 23 | $4NH_4^+ + 3HCO_3^- \rightarrow 4NO_2^- + 3CH_4 + 5H^+ + H_2O$ |
| E7 | 24 | $NH_4^+ + HCO_3^- \rightarrow NO_3^- + CH_4 + H^+$ |
| E11 | 25 | $4N_2 + 3HCO_3^- + 7H_2O \rightarrow 8NO_2^- + 3CH_4 + 5H^+$ |
| E12 | 26 | $4N_2 + 5HCO_3^- + 9H_2O \rightarrow 8NO_3^- + 5CH_4 + 3H^+$ |
| E15 | 27 | $4NO_2^- + HCO_3^- + H^+ + H_2O \rightarrow 4NO_3^- + CH_4$ |
| E19 | 28 | $H_2S + HCO_3^- + H_2O \rightarrow SO_4^{2-} + CH_4 + H^+$ |
| E26 | 29 | $4FeS_2 + 7HCO_3^- + 11H_2O \rightarrow 8SO_4^{2-} + 4Fe^{2+} + 7CH_4 + H^+$ |
| **Oxidation of $NH_4^+$** | | |
| F1 | 30 | $2NH_4^+ \rightarrow N_2 + 3H_2 + 2H^+$ |
| **Reduction of $N_2$** | | |
| G1 | 31 | $2N_2 + 6H_2O + 4H^+ \rightarrow 4NH_4^+ + 3O_2$ |
| G2 | 32 | $3H_2 + N_2 + 2H^+ \rightarrow 2NH_4^+$ |
| G4 | 33 | $3CH_4 + 4N_2 + 8H^+ + 6H_2O \rightarrow 8NH_4^+ + 3CO_2$ |
| G5 | 34 | $3CH_4 + 4N_2 + 5H^+ + 9H_2O \rightarrow 8NH_4^+ + 3HCO_3^-$ |
| G8 | 35 | $6H_2S + N_2 + 3Fe^{2+} \rightarrow 3FeS_2 + 2NH_4^+ + 4H^+$ |
| G10 | 36 | $3H_2S + 4N_2 + 2H^+ + 12H_2O \rightarrow 3SO_4^{2-} + 8NH_4^+$ |
| G13 | 37 | $3FeS_2 + 7N_2 + 8H^+ + 24H_2O \rightarrow 6SO_4^{2-} + 3Fe^{2+} + 14NH_4^+$ |

| | | |
|---|---|---|
| **Reduction of $NO_2^-$** | | |
| H1 | 38 | $4NO_2^- + 4H^+ \rightarrow 2N_2 + 3O_2 + 2H_2O$ |
| H2 | 39 | $2NO_2^- + 4H^+ + 2H_2O \rightarrow 2NH_4^+ + 3O_2$ |
| H3 | 40 | $3H_2 + NO_2^- + 2H^+ \rightarrow NH_4^+ + 2H_2O$ |
| H4 | 41 | $3H_2 + 2NO_2^- + 2H^+ \rightarrow N_2 + 4H_2O$ |
| H6 | 42 | $3CH_4 + 4NO_2^- + 8H^+ \rightarrow 4NH_4^+ + 3CO_2 + 2H_2O$ |
| H7 | 43 | $3CH_4 + 4NO_2^- + 5H^+ + H_2O \rightarrow 4NH_4^+ + 3HCO_3^-$ |
| H9 | 44 | $3CH_4 + 8NO_2^- + 8H^+ \rightarrow 4N_2 + 3CO_2 + 10H_2O$ |
| H10 | 45 | $3CH_4 + 8NO_2^- + 5H^+ \rightarrow 4N_2 + 3HCO_3^- + 7H_2O$ |
| H15 | 46 | $NH_4^+ + NO_2^- \rightarrow N_2 + 2H_2O$ |
| H18 | 47 | $3H_2S + 4NO_2^- + 2H^+ + 4H_2O \rightarrow 3SO_4^{2-} + 4NH_4^+$ |
| H21 | 48 | $3H_2S + 8NO_2^- + 2H^+ \rightarrow 3SO_4^{2-} + 4N_2 + 4H_2O$ |
| H25 | 49 | $3FeS_2 + 7NO_2^- + 8H^+ + 10H_2O \rightarrow 6SO_4^{2-} + 3Fe^{2+} + 7NH_4^+$ |
| **Reduction of $NO_3^-$** | | |
| I1 | 50 | $4NO_3^- + 4H^+ \rightarrow 2N_2 + 5O_2 + 2H_2O$ |
| I3 | 51 | $NO_3^- + 2H^+ + H_2O \rightarrow NH_4^+ + 2O_2$ |
| I4 | 52 | $4H_2 + NO_3^- + 2H^+ \rightarrow NH_4^+ + 3H_2O$ |
| I5 | 53 | $5H_2 + 2NO_3^- + 2H^+ \rightarrow N_2 + 6H_2O$ |
| I6 | 54 | $H_2 + NO_3^- \rightarrow NO_2^- + H_2O$ |
| I8 | 55 | $CH_4 + NO_3^- + 2H^+ \rightarrow NH_4^+ + CO_2 + H_2O$ |
| I9 | 56 | $CH_4 + NO_3^- + H^+ \rightarrow NH_4^+ + HCO_3^-$ |
| I11 | 57 | $5CH_4 + 8NO_3^- + 8H^+ \rightarrow 4N_2 + 5CO_2 + 14H_2O$ |
| I12 | 58 | $5CH_4 + 8NO_3^- + 3H^+ \rightarrow 4N_2 + 5HCO_3^- + 9H_2O$ |
| I14 | 59 | $CH_4 + 4NO_3^- \rightarrow 4NO_2^- + CO_2 + 2H_2O$ |
| I15 | 60 | $CH_4 + 4NO_3^- \rightarrow 4NO_2^- + HCO_3^- + H_2O + H^+$ |
| I22 | 61 | $5NH_4^+ + 3NO_3^- \rightarrow 4N_2 + 2H^+ + 9H_2O$ |
| I23 | 62 | $NH_4^+ + 3NO_3^- \rightarrow 4NO_2^- + 2H^+ + H_2O$ |
| I24 | 63 | $N_2 + 3NO_3^- + H_2O \rightarrow 5NO_2^- + 2H^+$ |
| I25 | 64 | $8H_2S + NO_3^- + 4Fe^{2+} \rightarrow 4FeS_2 + NH_4^+ + 6H^+ + 3H_2O$ |
| I27 | 65 | $H_2S + NO_3^- + H_2O \rightarrow SO_4^{2-} + NH_4^+$ |
| I28 | 66 | $10H_2S + 2NO_3^- + 5Fe^{2+} \rightarrow 5FeS_2 + N_2 + 8H^+ + 6H_2O$ |
| I30 | 67 | $5H_2S + 8NO_3^- \rightarrow 5SO_4^{2-} + 4N_2 + 2H^+ + 4H_2O$ |
| I31 | 68 | $2H_2S + NO_3^- + Fe^{2+} \rightarrow FeS_2 + NO_2^- + 2H^+ + H_2O$ |
| I33 | 69 | $H_2S + 4NO_3^- \rightarrow SO_4^{2-} + 4NO_2^- + 2H^+$ |
| I38 | 70 | $4FeS_2 + 7NO_3^- + 6H^+ + 11H_2O \rightarrow 8SO_4^{2-} + 4Fe^{2+} + 7NH_4^+$ |
| **Oxidation of Sulfide with Fe** | | |
| J1 | 71 | $2H_2S + Fe^{2+} \rightarrow H_2 + FeS_2 + 2H^+$ |
| **Reduction of $SO_4^{2-}$** | | |
| M1 | 72 | $SO_4^{2-} + 2H^+ \rightarrow H_2S + 2O_2$ |
| M2 | 73 | $4H_2 + SO_4^{2-} + 2H^+ \rightarrow H_2S + 4H_2O$ |
| M3 | 74 | $7H_2 + 2SO_4^{2-} + Fe^{2+} + 2H^+ \rightarrow FeS_2 + 8H_2O$ |
| M6 | 75 | $CH_4 + SO_4^{2-} + 2H^+ \rightarrow H_2S + CO_2 + 2H_2O$ |
| M7 | 76 | $CH_4 + SO_4^{2-} + H^+ \rightarrow H_2S + HCO_3^- + H_2O$ |

| | | |
|---|---|---|
| M9 | 77 | $7CH_4 + 8SO_4^{2-} + 4Fe^{2+} + 8H^+ \rightarrow 7CO_2 + 4FeS_2 + 18H_2O$ |
| M10 | 78 | $7CH_4 + 8SO_4^{2-} + 4Fe^{2+} + H^+ \rightarrow 7HCO_3^- + 4FeS_2 + 11H_2O$ |
| M21 | 79 | $4NH_4^+ + 3SO_4^{2-} \rightarrow 4NO_2^- + 3H_2S + 2H^+ + 4H_2O$ |
| M22 | 80 | $NH_4^+ + SO_4^{2-} \rightarrow NO_3^- + H_2S + H_2O$ |
| M29 | 81 | $4N_2 + 3SO_4^{2-} + 4H_2O \rightarrow 8NO_2^- + 3H_2S + 2H^+$ |
| M30 | 82 | $4N_2 + 5SO_4^{2-} + 2H^+ + 4H_2O \rightarrow 8NO_3^- + 5H_2S$ |
| M35 | 83 | $4NO_2^- + SO_4^{2-} + 2H^+ \rightarrow 4NO_3^- + H_2S$ |
| **Oxidation of $H_2S$** | | |
| M1a | 84 | $H_2S + 2O_2 \rightarrow SO_4^{2-} + 2H^+$ |
| M6a | 85 | $H_2S + CO_2 + 2H_2O \rightarrow CH_4 + SO_4^{2-} + 2H^+$ |

**Supplemental Table 3. Endmember hydrothermal fluid compositions used in the energetics calculations**

| | Strytan | Strytan model | Eridania (5% Ol) | Eridania (16% Ol) |
|---|---|---|---|---|
| **pH** | 10.03 | 9.79 | 7.96 | 7.65 |
| **$CH_4$** | $5.12 \times 10^{-7}$ | *(ab)* | $1.60 \times 10^{-2}$ | $6.66 \times 10^{-3}$ |
| **$CO_{2\,(aq)}$** | *n/a* | $3.33 \times 10^{-8}$ | $1.09 \times 10^{-8}$ | $1.80 \times 10^{-6}$ |
| **$H_2$** | $1.44 \times 10^{-3}$ | $1.44 \times 10^{-3}$ | $7.70 \times 10^{-7}$ | $4.93 \times 10^{-7}$ |
| **$H_2S$** | $1.25 \times 10^{-5}$ | $1.25 \times 10^{-5}$ | $8.95 \times 10^{-11}$ | $2.30 \times 10^{-10}$ |
| **$HCO_3^-$** | *n/a* | $1.37 \times 10^{-4}$ | $8.67 \times 10^{-5}$ | $8.00 \times 10^{-5}$ |
| **$N_{2\,(aq)}$** | *n/a* | *n/a* | $5.21 \times 10^{-2}$ | $4.90 \times 10^{-2}$ |
| **$NH_4^+$** | $1.55 \times 10^{-6}$ | *n/a* | 0.496 | 0.660 |
| **$NO_2^-$** | $4.34 \times 10^{-8}$ | *n/a* | *(ab)* | *(ab)* |
| **$NO_3$** | $7.26 \times 10^{-7}$ | *n/a* | *(ab)* | *(ab)* |
| **$O_{2\,(aq)}$** | *n/a* | *(ab)* | *(ab)* | *(ab)* |
| **$SO_4^{2-}$** | $2.97 \times 10^{-7}$ | $4.12 \times 10^{-7}$ | *(ab)* | *(ab)* |
| **$Fe^{2+}$** | $1.48 \times 10^{-7}$ | $3.22 \times 10^{-12}$ | $4.23 \times 10^{-8}$ | $2.26 \times 10^{-7}$ |

Units are molal (mol/kg)

*(ab)*: concentrations lower than $10^{-12}$ and therefore considered to be absent from the fluid

n/a: concentrations not measured

[a] values from Kristjansdottir, unpublished
